# Soil iron drives beneficial maize microbiome feedbacks in rotations with wheat

**DOI:** 10.64898/2026.01.08.698436

**Authors:** Jan Waelchli, Henry Janse van Rensburg, Katja Stengele, Viola D’Adda, Selma Cadot, Veronica Caggìa, Valentin Gfeller, Klaus Schlaeppi

## Abstract

**Background:** Plants change their surrounding soil microbiome by root exudates and such conditioned microbiomes impact the performance of the present as well as the next plant generation as for example in crop rotations. The big challenge is that such ‘microbiome feedbacks’ are highly context-dependent, i.e. they vary in strength and direction dependent on the local soil environment – of which the driving factor(s) remain unknown. Including maize in crop rotations involves benzoxazinoids (BXs), which are exuded from roots and alter the soil microbiome, which in turn affects growth and defence of the following crop.

**Results:** Here, we grew wild-type and BX-depleted maize in the field to differentially condition their soil microbiome and we found varying feedbacks on wheat performance dependent on the local physicochemical soil parameters. Using multivariate, correlation and modelling approaches and including additional data from two previous field experiments, we identified plant-available (PA) iron to explain BX-dependent microbiome feedbacks on wheat. The BX-conditioned soil microbiome caused wheat to grow taller at low levels of soil PA-iron but smaller at high levels. This finding was generalized testing these maize microbiome feedbacks on the model plant *Arabidopsis thaliana* using soil batches containing different levels of iron. Consistent with wheat, a significant inverse relationship between soil PA-iron levels and plant growth was found. This relationship was experimentally validated with *Arabidopsis* grown at low levels of soil iron where iron supplementation abolished the beneficial feedback of the BX-conditioned soil microbiome.

**Conclusion:** Together these findings revealed that beneficial microbiome feedbacks occur at low levels of plant-available iron, i.e. when plants grow in a suboptimal soil, but they are lost when plants are nutritionally well supported. These results underscore the importance of iron availability in soil for beneficial microbial feedbacks on plant growth and predict agronomic benefits of incorporating maize in crop rotations on low iron soils.

## Introduction

Plants condition their local soil environment by the release of root exudates (Sasse et al., 2018). This form of soil conditioning, modifying the physicochemical as well as microbial properties, often influences the performance of subsequent plants growing in the same soil – a process referred to as plant-soil feedbacks (PSF; Bezemer et al., 2006; van der Putten et al., 2016). If an altered soil microbiome is the proximal cause for altered performance, then these plant responses are referred to as ‘microbiome feedbacks’ (Janse van Rensburg et al., 2024). Such soil or microbiome feedbacks on plant growth, defence or development can have profound implications in practice (e.g., crop yields) or nature (e.g., ecosystem succession dynamics; Mariotte et al., 2018). In agriculture, such soil or microbiome feedbacks operate in crop rotations where the past crops impact the yield of the subsequent crop. Today, optimal rotation schemes take advantage of feedbacks that enhance soil fertility, promote beneficial microbial communities, suppress soil-borne pathogens and reduce plant pests (Gfeller et al., 2023b; Kirkegaard et al., 2008; Peralta et al., 2018). Altering or co-cultivating crops started with Neolithic farming in the Near East (Zohary and Hopf, 1973) and more optimal rotation schemes continue to develop with continuous learning (Mudare et al., 2025). While farmers have learned and identified cropping sequences that exploit positive over negative soil or microbiome feedbacks, the underlying mechanisms often remain unknown today.

Plant-soil and microbiome feedbacks are highly context-dependent. Feedbacks vary not only in strength but also in their direction (they can be negative or positive) depending on the soil environment. This high context-dependency is thought to result from the interplay between the soil microbiota and the abiotic soil properties, including physical (e.g., structure, moisture) and chemical (e.g., nutrient availability, plant exudates) parameters (Cheng et al., 2024). Typically, feedback responses are not fully consistent *between* different fields (Aaronson et al., 2023), which is not surprising as different soil types differ in many ways such as in their physicochemical parameters. Feedback responses vary also in a more fine-grained manner due to local soil heterogeneity *within* fields. Farmers experience such context-dependency of microbiome feedbacks when observing over– and under-yielding areas in their fields (Maestrini and Basso, 2018). Deciphering the context-dependency of strength and direction is of key importance towards understanding the underlying mechanisms of microbiome feedbacks.

Maize (*Zea mays* L.) and wheat (*Triticum aestivum* L.) are two of the most commonly grown crops worldwide (Vasileiadis et al., 2017). Both crops are typically cultivated based on heavy use of agrochemicals, responsible for negative environmental impacts (Meissle et al., 2010; Pimentel, 2005), and therefore, in focus for more sustainable agricultural practices. Among these, maize-wheat rotations for instance become more widely practiced in Europe (Meissle et al., 2010). Like other sweet grasses, maize and wheat produce benzoxazinoids (BXs), a class of multifunctional plant secondary metabolites (Niemeyer, 2009). BXs are defence compounds against generalist insects, have antimicrobial activity, they function allelopathically as phytotoxins, serve as phytosiderophores for iron uptake or they alleviate abiotic stress by soil aluminium and arsenic (Zhou et al., 2018). Importantly, BXs are exuded from the roots to the surrounding soil, where they strongly alter the microbial communities (Cadot et al., 2021b; Cotton et al., 2019; Hu et al., 2018b; Kudjordjie et al., 2019). Maize secretes quantitatively more BXs than wheat and predominantly DIMBOA, DIMBOA-Glu and HDMBOA-Glu (Hu et al., 2018; see methods for full chemical names). Thus, sweet grasses like maize and wheat condition the microbiomes in the surrounding soil by exudation of BXs.

BX-conditioned soil microbiomes (referred to as ‘BX_plus_ microbiomes’) drive differential feedbacks on the next plant generation as observed in experimental maize-maize (Hu et al., 2018) and maize-wheat (Cadot et al., 2021a) rotations in the greenhouse. These conclusions are derived from comparing plant performance on BX_plus_ relative to the control ‘BX_minus_ soil’ (i.e., microbiome conditioned by BX-defective maize plants). For instance, wheat grew smaller but was more resistant to insects in response to the BX_plus_ microbiome (Cadot et al., 2021a). Feedbacks driven by BX_plus_ microbiomes were also found for the model plant *Arabidopsis thaliana* (hereafter Arabidopsis) growing in maize conditioned soil in climate chambers (Stengele et al., 2024). The reference accession Columbia-0 grew larger, was developmentally more advanced and was more resistant to the fungal pathogen *Botrytis cinerea*. Furthermore, Arabidopsis revealed massive genetic variation in their growth feedbacks, allowing the identification of a plant immune receptor mediating these microbiome feedbacks on plant growth (Janse van Rensburg et al., 2025). All above reported work on feedbacks of maize, wheat or Arabidopsis in response to BX_plus_ microbiomes was performed in controlled conditions using soil from the same field site (Changins, Switzerland). For the many experiments we repeatedly went back to this site and collected batches of soil from different spots of the field. Over the many experiments, we noticed strong variation in the strength of the feedbacks between soil batches and initially, dismissed these observations as experiment– to-experiment variations [Note: this study here now reveals that the variation in feedback strength is due to local soil heterogeneity from where the different soil batches were collected].

Complementary to the work in controlled conditions (paragraph above), we have tested the agronomic relevance of the feedbacks of maize BX_plus_ vs. BX_minus_ microbiomes on wheat in agronomically realistic conditions at two field sites (Gfeller et al., 2023b, 2023a). In the first season, we cultivated wild-type BX producing and BX-deficient maize lines to differentially condition the soil and then assessed in the following season the feedbacks of BX_plus_ vs. BX_minus_ microbiomes on winter wheat. In the first field site (Posieux), we found that soil conditioning by BXs increased wheat emergence, tillering, height, and biomass and decreased insect herbivore abundance at the same time (Gfeller et al., 2023b). Importantly, these BX-induced microbiome feedbacks increased wheat yield by 4% without a reduction in grain quality in this maize-wheat rotation. These BX-induced microbiome feedbacks were confirmed in a repeat experiment at the second field site (Changins, same site as for soil collection) – but they were not apparent at first glance (Gfeller et al., 2023a). Averaged across the field, we did not detect field-scale feedbacks on wheat performance. However, taking the local soil condition into account, significant feedbacks were found along the physicochemical gradient existing in that field. For instance, wheat gradually responded to the BX_plus_ microbiome with varying heights along the soil gradient; i.e., BX_plus_ microbiome caused increased height at one and decreased height at the opposing end of the field gradient. Taken together, BX conditioned soil microbiomes drive feedbacks on wheat in agronomically realistic conditions, but they vary in strength and direction depending on local soil physicochemical heterogeneity.

Here in this study, we tested the reproducibility of the aforementioned findings that soil conditioning by BX-producing plants increased wheat growth (Gfeller et al., 2023b) and that these microbial feedbacks vary in strength and direction depending on soil heterogeneity (Gfeller et al., 2023a). Analogous to our two previous field studies, we conducted a third experiment at a new field site growing wild-type and BX-deficient maize to differentially condition the soil microbiomes and using wheat to score feedbacks (**Fig. 1AB**). We captured local soil heterogeneity in each replicated plot by measuring the physicochemical parameters and the microbial communities, which maize plants leave as legacy for wheat growth. Through multivariate, correlation and modelling analyses and combining the data from the three field experiments, we confirmed the differing microbial communities due to BX conditioning and that the strength of their feedbacks on wheat largely depended on the availability of soil iron levels. We extended our analysis to experiments with Arabidopsis where the strength of the feedbacks varied by soil batches. Like wheat, Arabidopsis showed an inverse relationship between soil iron and microbial feedbacks, which we could experimentally validate with iron supplementation. Overall, our work revealed that low levels of soil iron are needed so that the BX conditioned soil microbiomes promote growth as feedbacks on the subsequent plant generation.

**Figure 1.**
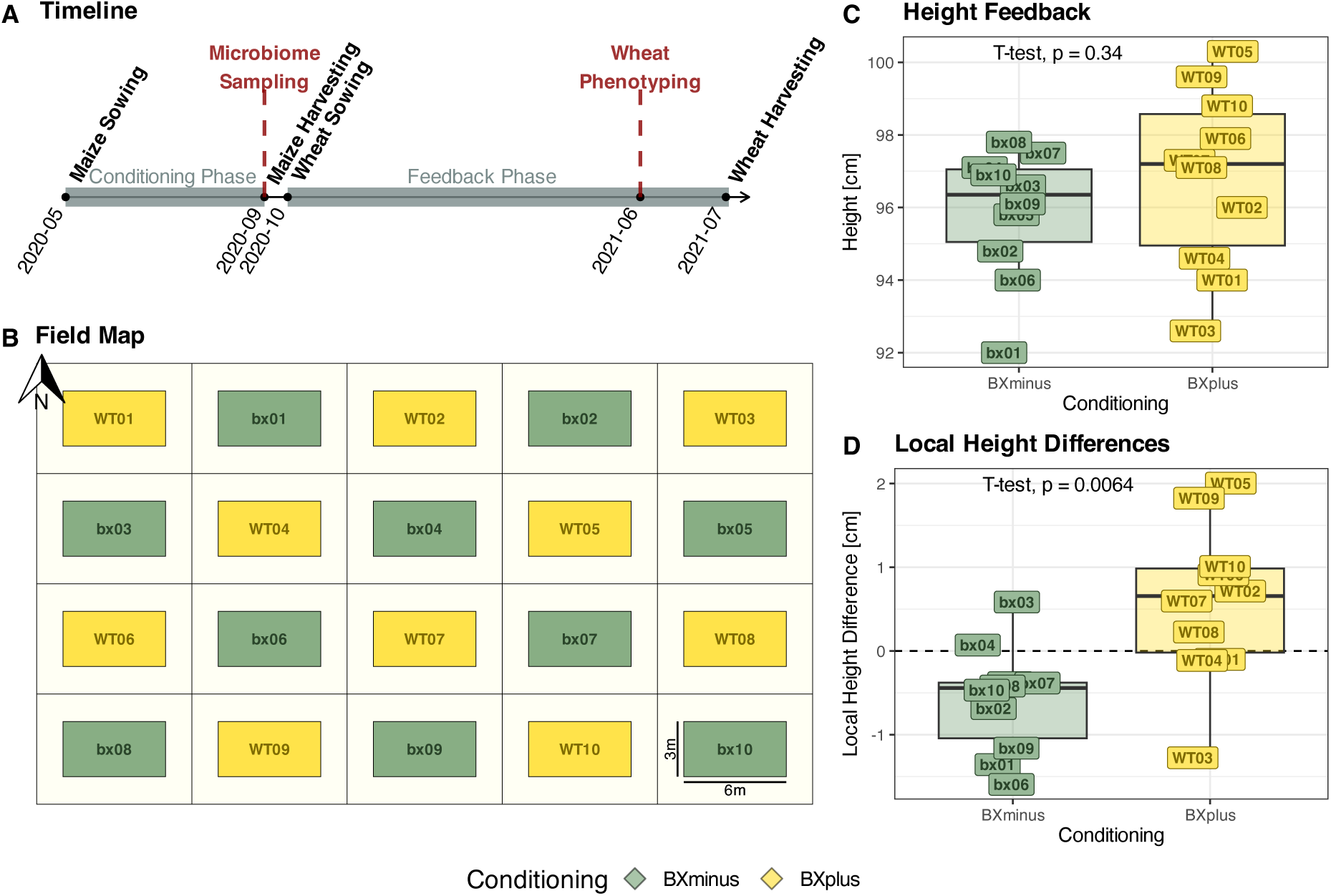
| Maize-wheat plant-soil feedback experiment in Reckenholz. **(A)** At the Reckenholz field site a maize-wheat feedback experiment was performed. **(B)** A total of 20 3 × 6 m plots were established, half of it were conditioned with wild-type maize (BX_plus_ conditioning) and the other half with the benzoxazinoid-defective bx1 mutant (BX_minus_ conditioning). Each plot was surrounded by a 3 m-wide buffer zone planted with *phacelia*. **(A)** At the end of the conditioning phase, the maize root microbiome was sampled, physicochemical soil properties were measured and maize was harvested. Following the conditioning phase, wheat was grown on the conditioned soils and wheat phenotypes were measured towards the end of the feedback phase. The detailed phenotyping of wheat is described in **Fig. S2** while here, plant height **(C)** is shown as an example. Wheat did not differ in height whether grown on BX_plus_ or BX-conditioned soils (t-test, p-value displayed in panel). **(D)** However, taking spatial field heterogeneity into account, i.e. calculating the “local height differences” by normalizing each plot to its neighboring plots (see Methods), wheats differ in height between BX_plus_ and BX_minus_ conditioned soils (t-test, p-value displayed in panel).

## Material & Methods

### Maize-Wheat rotation experiment in the field

The setups of the field experiments in Changins and Posieux have been described earlier in Gfeller *et al*. (2023a) and (2023b), respectively. Here we document the specifications, plant material, setup and sampling of the new field experiment performed at Reckenholz (Zürich, Switzerland). The field is called “Schlag 109” (47°25’45’N, 8°31’23’E) and belongs to the research station Agroscope. This site has a loamy soil (clay ∼21%, silt ∼31% and sand ∼46%) with pH 6.8 and a humus content of 3%. The more detailed physicochemical soil analysis is documented on GitHub (dataset in section **B01**). The field’s history was three years of fodder meadow (mixture of red clover and Italian ryegrass) and it was managed according to Swiss conventional agricultural practices by the field team of Agroscope.

The experiment consisted of a maize-wheat rotation with maize cultivated during the conditioning phase in 2020 and wheat in 2021 in the feedback phase (**Fig. 1A**). For soil conditioning, we grew wild-type maize (*Zea mays*; inbred line W22) and the benzoxazinoid-deficient mutant *bx1* in W22 background (Tzin et al., 2015). After clearing the preceding meadow with Roundup (8 L/ha) in autumn 2019, the seed bed was prepared in spring 2020 with a harrow, weeds controlled with herbicides (Aspect 1.5 L/ha, Banvel 4S 0.5 L/ha and Laudis 2 L/ha) and the seeds then directly sown with a seed drill (seeding: 28.05.2020). In total, we used 2’500 seeds wild-type and 2’750 seeds the *bx1* mutant maize to correct for the lower germination rate of the latter (determined ahead). Maize was sown in rows with 75 cm distance between rows. Wild-type and *bx1* mutant maize were alternately seeded with 10 replicate plots each following a chessboard design (**Fig. 1B**). Each plot measured 6 x 3 m ‘hosting’ 4 rows of maize (**Fig. S1A**) and was surrounded by a 3 m buffer zone, where we planted *Phacelia*. The field was minerally fertilised with a total of 115 kg/ ha N in form of Urea applied in two doses. Harvest was after 14 weeks (harvest: 08.09.2020): At two locations per plot, we selected single maize plants of which we sampled the underneath soil (cores of 20 x 20 x 20 cm) and root systems (**Fig. S1A**). Both the soil and root samples were mixed per plot for subsequent analyses of physicochemical soil parameters and root microbiota profiles, respectively, as described below.

For the feedback phase (**Fig. 1A, Fig. S1B**), we then cultivated the winter wheat variety ‘CH Claro’ (referred as Claro; UFA Samen, www.ufasamen.ch; same variety as in Changins and Posieux). Seed bed preparation included a ploughing and harrowing of the field followed by direct sowing with a seed drill (seeding: 22.10.2020, 1.6 dt/ha). The wheat was sown in rows with 15 cm distance between the rows across all plots. The field was minerally fertilised with three applications of ammonium nitrate (1: 48 kg/ha of 24% N, 5% Mg, 6% S; 2: 60.5 kg/ha 27.5% N and 3: 24 kg/ha 24% N, 5% Mg, 6% S) equivalent to a total of 132.5 kg/ ha N and the following plant protection products were applied: herbicides Globus 0.07 L/ha and Othello 1 L/ha and the insecticide Audienz 0.05 L/ha. On 11.06.2021, when all plants have reached the stage of fruit developing (BBCH 70-79), we selected in 3 lines 10 random wheat plants within a 4 m segment of the plots to get the mean plant height (cm), mean chlorophyll content (determined with a Minolta SPAD meter) and mean insect infestation by counting the number of larvae from the *Oulema melanopus* beetle on the two newest tillers. The aboveground biomass of 1 m of a wheat row was sampled (**Fig. S1B, Fig. S2ABC**), dried in the oven at 60° C for 48h and the dry weight was determined (g/m). After this, we re-established the plot outlines by mowing to keep a 4 x 1.5 m core-zone based on GPS coordinates. We finally harvested on 30.07.2021 the core zone and determined the kernel yield (g/m^2^) (**Fig. S1B, Fig. S2D,** dataset on GitHub, section **B04**).

### Soil physicochemical analysis

The collection of soil cores was described above. Per plot 500 g of soil was sent to LBU Laboratories (Eric Schweizer AG, Thun; Switzerland) for analysis of physicochemical soil parameters. The same parameters as in Gfeller et al. (2023a, 2023b) were measured. Five plots could not be analysed due to a lack of soil (BX2, BX5, WT2, WT3, WT5). Water (H₂O) extractions estimate plant-available (PA) nutrients, carbon dioxide (CO₂) extractions measure partly plant-available nutrients (PPA), and ethylenediaminetetraacetic acid (EDTA) extractions determine nutrients that are present in the plant-unavailable (UA) fraction (dataset on GitHub, section **B01**).

### Maize root microbiome analysis

The collection of soil cores was described above. To prepare root microbiota samples we followed Gfeller et al. (2023b) with the following changes: Maize samples were stored at room temperature and immediately processed and wheat samples were stored at –20°C. Wheat samples were lyophilized for 72h. Root fragments were ground in 25 mL grinding jars with a 1.5 cm steel ball using a ball mill (Retsch, Germany) for 120 s by 30 Hz. Samples were placed in the double amount of extraction buffer for 10 minutes. 125 mg of root powder was used for DNA extraction. DNA extraction was done with the KF MagAttract Power Soil Kit (Qiagen, Germany) on a KingFisher robot (Thermo Fisher, USA) and DNA concentrations were measured with the AccuClear Kit (Biotium, USA). In the first PCR, the DNA template was diluted to a final concentration of 1 ng per reaction. After library preparation, samples were sequenced using a paired-end 300 bp v3 chemistry on an Illumina MiSeq instrument at the NGS platform of the University of Bern (ngs.unibe.ch). The raw sequencing data is made available at ENA (see below).

### Maize-Wheat feedback experiments

All analyses of the maize-wheat experiments were conducted for the field in Reckenholz alone and also combined with the data from the previous field experiments in Changins (Gfeller et al., 2023a) and Posieux (Gfeller et al., 2023b).

#### Root Microbiome

Root microbiome raw sequencing (available on ENA, see below) data from Changins, Posieux and Reckenholz were combined. Then we followed the same pipeline as described in Gfeller et al. (2023b) to infer exact amplicon sequencing variants (ASVs) and assign taxonomies. These calculations were performed at sciCORE (http://scicore.unibas.ch/), the scientific computing centre of the University of Basel (metafile and code are on **GitHub A**, the processed sequencing dataset in section **B02**). We used the R package vegan (version 2.6.8, Oksanen et al., 2024), followed the recommendation of Weiss et al. (2017) and normalized the samples by rarefying at thresholds of 8’000 and 800 bacterial and fungal sequences, respectively. Samples with less sequences and samples detected as outliers by the CLOUD method (nearest neighbours=40%, outlier percentile=0.1, Montassier *et al*., 2018) were discarded. In a next step, we defined a core microbiome containing ASVs with an occurrence in at least 10% of the samples and a minimal abundance of >0.01%. Community effects on beta diversity were tested with permutational analysis of variance (PERMANOVA, 999 permutations) on Bray-Curtis distances. For Reckenholz plots, we tested for differences in conditioning and visualized it by the first two axes of a Principal Coordinates Analysis (PCoA).

Taking all field sites together, we investigated for differences in conditioning, field and conditioning-field interaction and showed the results by plotting the first two axes of a Canonical Analysis of Principal coordinates (CAP, package phyloseq 1.48.0; McMurdie and Holmes, 2013). Dataset, code and analysis documentation are provided on GitHub, section **B02**.

#### Physicochemical soil parameters

We investigated for differences in conditioning, soil and their interaction using a PERMANOVA (999 permutations) on Euclidean distances and displayed results in a Principal Component Analysis (PCA), and as biplot with the package FactoMineR (version 2.11, Lê et al., 2008), and the package factoextra (version 1.0.7, Kassambara and Mundt, 2020). Dataset, code and analysis documentation are provided on GitHub, section **B01**.

#### Benzoxazinoid analysis

For statistical analysis we used 2,4-dihydroxy-7-methoxy-1,4-benzoxazin-3-one (DIMBOA), 2-O-β-D-glucopyranosyl-2,4-dihydroxy-7-methoxy-(2H)-1,4-benzoxazin-3(4H)-one (DIMBOA-Glucose) and 2-hydroxy-4,7-dimethoxybenzoxazin-3-one (HDMBOA-Glucose) measurements from Changins and Posieux sites measured on BX_plus_ conditioned soil at the end of the conditioning phase. Methods were described in Gfeller et al. (2023a) and (2023b). No benzoxazinoids were measured in Reckenholz. For details see GitHub, section **B03**.

#### Plant feedbacks

The variation in plant height between field sites was normalized by z-score normalization per field. Following Gfeller *et al*. (2023a), we calculated local height differences by dividing plant heights of each BX_plus_ plot by the mean of plant heights from the surrounding BX_minus_ plots and vice versa. With a t-test, we investigated for height differences between conditionings (GitHub, section **B04**). Further, we inversed the local height differences of BX_minus_ plots (multiplied by –1) to calculate the local height feedback as the absolute difference between neighbouring plots (BX_plus_ – BX_minus_). Positive feedback: plants on BX_plus_ conditioned plots grow higher than plants on BX_minus_ conditioned plots. Negative feedback: plants on BX_plus_ conditioned plots grow smaller than on BX_minus_ plots. We correlated local height feedbacks with the PCA axes from the physicochemical soil analysis, by performing a Pearson correlation test. Local height feedback from plots without physicochemical soil measurements were excluded. Mean local height feedback were used for plots where soil was pooled for physicochemical soil parameters (GitHub, section **B05**). Local height feedback was also correlated to the PCoA axes of bacteria and fungi. Samples without bacterial or fungal root microbiome measurements and samples which were discarded during microbial analysis were removed (GitHub, section **B06**).

#### Modelling

Samples without measurements of physicochemical soil parameters, bacterial abundance or fungal abundance were removed. As described above, samples were pooled, whenever physicochemical soil parameters were measured between plots. Due to the high dimensionality of our dataset (more explanatory variables than samples) and the presence of substantial multicollinearity among variables, we used Least Absolute Shrinkage and Selection Operator (LASSO) regression (package glmnet 4.1.8, Friedman *et al*., 2023) to select the best prediction model (Friedman et al., 2010). As explanatory variables, we included all measured physicochemical soil parameters and all bacterial and fungal ASVs that occurred in at least 4 samples and had a relative abundance of at least 5% of the total sequence count. Using k-fold cross-validation, we determined the optimal value of the regularization parameter λ (L1 penalty). We selected λ such that the model error remained within one standard error of the minimum cross-validation error. Variables with non-zero coefficients were retained as predictors.

Given that model selection can be unstable in the presence of high multicollinearity and sparsity among predictors, as numerous zero values, we did a repeated-subsampling stability analysis to assess the robustness of variable selection and mitigate seed dependency (Heinze et al., 2018). Inspired by the approach of Meinshausen and Bühlmann (2010), we ran the model 500 times using 80% of the samples in each iteration, without replacement. While re-estimating λ in each iteration prevents formal control of the expected number of false positives, we report the frequency of variable selection across iterations as an indicator of stability. Predictors selected in less than 25% of iterations were considered unstable and were excluded. The remaining predictors were categorized based on selection stability as low (25– 50%), moderate (50–75%), or high (75–100%). Finally, we evaluated the adjusted R² and p-value of the final model using Analysis of Variance (ANOVA), and calculated adjusted relative importance values for each retained predictor by averaging over predictor orderings (package relaimpo 2.2.7, Groemping, 2023; see GitHub, section **B07**).

### Maize-Arabidopsis feedback experiments

Multiple independent maize – Arabidopsis feedback experiments were performed. We combined their data to investigate them collectively. Details on GitHub, section **B08**.

#### Experimental setups

For each experiment, field soil was conditioned by growing wild-type maize and *bx1* mutants for 12 weeks. Conditioning was either carried out directly in the field or soil was previously collected, sieved (1 cm mesh) and maize was grown in individual pots in the greenhouse or Phytotron chambers. For details see **Table S3**, Stengele *et al*. (2024) and Janse van Rensburg *et al*. (2025). At the end of the conditioning phase, conditioned BX_minus_ and BX_plus_ soils from repeat field plots or pots were pooled by maize genotype and experiment. A 500 g sample of each conditioned soil from each soil batch was sent to LBU Laboratories (Eric Schweizer AG, Thun; Switzerland) for analysis of physicochemical soil parameters. The same parameters as for the previously described Reckenholz samples were analysed. After the conditioning phase, the soil was mixed with 20% autoclaved quartz sand and pots (5.5 x 5.5 x 5 cm) were filled by weight. The pots were sown with stratified seeds of the reference accession Columbia-0 (3 d at 4 °C). Plants were grown in Percival growth chambers (CLF Plant Climatics, Wertingen, Germany) with 10 h of light (100 μmol m^−2^ s^−1^, 21 °C, 60% relative humidity) and 14 hours of darkness (0 μmol m^−2^ s^−1^, 21 °C 18 °C, 60 % relative humidity). They were watered 3 times a week by weight and randomized once per week. Plants were either not fertilized, fertilized with 10 mL of 1:3 times diluted ½ strength Hoagland solution at 21 d or 14 d and 21 d after germination or fertilized with 5 ml of undiluted ½ strength Hoagland solution 21 d and 35 d after germination. Plants were photographed 40±2 d after germination. Detailed information for each experiment is given in **Table S3**.

#### Feedback analyses

To determine Arabidopsis growth, plant-photographs were processed using the ARADEEPOPSIS 1.3.1 software (Hüther and Schandry, 2021) to compute the rosette area for each sample. Rosette areas were normalized using z-score normalization within each experiment. Similar to local height differences used in the wheat experiments, we calculated rosette differences for Arabidopsis. For each data point *x* within an experiment *e*, the rosette differences was calculated as 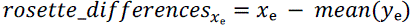, where *y*e represents all other data points in the same experiment but of the other conditioning. Then we calculated the rosette feedback by multiplying BX_minus_ conditioned rosette differences by –1. Rosette feedbacks were correlated to PA-iron by performing a Pearson correlation test. To investigate for differences in the correlation coefficients between PA-iron local rosette feedback correlation and PA-iron local height feedback correlation, a Fisher test for independent groups was performed (package cocor 1.1.4, Diedenhofen, 2022).

#### Iron supplementation experiment

Finally, we performed an iron supplementation experiment using the low-iron soil batch BS09. For the low iron treatment, we supplement soil during the conditioning phase with 100 mL of 0.2% Plantaaktiv Typ K (Hauert HBG Duenger AG, Grossaffoltern, Switzerland) and 0.001% Sequestrene Rapid (Maag, Westland Schweiz GmbH, Dielsdorf, Switzerland). For the high iron treatment, the first four weeks of conditioning was done identical to the low iron treatment before increasing the concentrations to 200 mL of 0.2% Plantaaktiv Typ K, 0.02% Sequestrene Rapid for the remaining 8 weeks of the conditioning phase. For both fertilization treatments, a t-test was used to test for differences in rosette size of Arabidopsis grown on BX_plus_ and BX_minus_ conditioned soil.

### Hardware and Software

If not explicitly mentioned, analyses were performed in R 4.5.1 (R Core Team, 2024) on a MacBook Pro 2020 with a Quad-Core Intel Core i7 processor. For data organisation and transformation, the tidyverse-package collection 2.0.0 (Wickham et al., 2019) and gtools 3.9.5 (Warnes et al., 2023) were used. Figures were created with ggplot2 3.5.2 from the tidyverse and the data-visualization packages cowplot 1.1.3 (Wilke, 2024), ggpubr 0.6.0 (Kassambara, 2023), gghx4 0.3.0 (Brand, 2024), ggnewscale 0.5.1 (Campitelli, 2024), scales 1.4.0 (Wickham et al., 2023), terra 1.8.54 (Hijmans, 2024), tidyterra 0.7.2 (Hernangómez, 2024) and geomtextpath 0.1.5 (Cameron and Brand, 2024). Tables were visualized by the package kableExtra 1.4.0 (Zhu, 2024).

## Data and code availability

The raw microbiome sequencing data is deposited at the European Nucleotide Archive (http://www.ebi.ac.uk/ena) under the study accessions PRJEB59165, PRJEB53704 and PRJEB94164. Datasets and code used in this study to analyse microbiome sequencing data, to do statistical analyses and to create figures are available on GitHub (https://github.com/PMI-Basel/Waelchli_et_al_Reckenholz_field_experiment). The GitHub repository has two parts with “A” containing information and bash and R code for microbiome analysis as follows: A1) Metafile containing barcode, primer and sample assignment information A2) raw sequencing file names and A3) code to analyse microbiome sequencing data. The part “B” contains the raw data and R code for the following analyses: B01) soil chemistry, B02) statistic microbiome analysis, B03) BX soil concentrations, B04) wheat feedback, B05) local height feedback – soil chemistry interaction, B06) local height feedback – microbiome interaction, B07) modelling approach, B08) Arabidopsis feedback experiments.

## Results

### Feedbacks on wheat depend on local soil condition

To assess the role of soil conditions on wheat feedbacks, we performed a third field trial (site: Reckenholz), where analogous to the two previous experiments (sites Posieux and Changins), we first cultivated wild-type maize (inbred line W22) and the benzoxazinoid-deficient mutant *bx1* (in W22 background) to differentially condition the soil (**Fig. 1AB**). Then, we assessed the feedbacks on the winter wheat variety Claro in response to BX_plus_ vs. BX_minus_ soil microbiomes in the following season. Similar to the heterogenous field in Changins (Gfeller et al., 2023a), field-scale feedbacks were not observed at Reckenholz while wheat performance differed significantly when considering the *local* feedback (takes local soil condition into account, see Methods). We exemplify this conclusion with wheat height (**Fig. 1CD**) and document the same for biomass, chlorophyll content, insect infestation and kernel yield in **Fig. S2**. Feedbacks on plant height revealed no difference between the two conditioned soils across the entire field (*p* > 0.05, **Fig. 1C**), whereas the *local* height feedback revealed wheat to grow better on BX_plus_ compared to BX_minus_ soils (*p* < 0.01, **Fig. 1D**). Thus, we found also for the Reckenholz site that soil conditioning by BX-producing plants increased wheat performance in a local soil context-dependent manner.

### Local physicochemical soil heterogeneity correlates with strength of feedbacks

Finding that wheat performance was affected by local soil condition (**Fig. 1**), the next step was to search the soil parameters that could explain the local differences in growth feedbacks. For this, we analysed in each plot the physicochemical parameters and the bacterial and fungal communities that maize plants leave as legacy for wheat growth. We used permutational analysis of variance (PERMANOVA) for quantification of effect sizes (approximated based on R^2^ values) and unconstrained ordination for visualisation. The physicochemical parameters did not differ between soils conditioned by wild-type or *bx1* maize plants (R^2^ = 1.8%, *p* > 0.05, **Fig. 2A**). In contrast, larger effects sizes and significant differences between wild-type and *bx1* maize plants were found in root bacterial (R^2^ = 20.7%, *p* < 0.05, **Fig. 2B**) and fungal communities (R^2^ = 13.0%, *p* < 0.05, **Fig. 2C**). Thus, the differential conditioning affected the microbial but not the physicochemical parameters of the soil.

**Figure 2.**
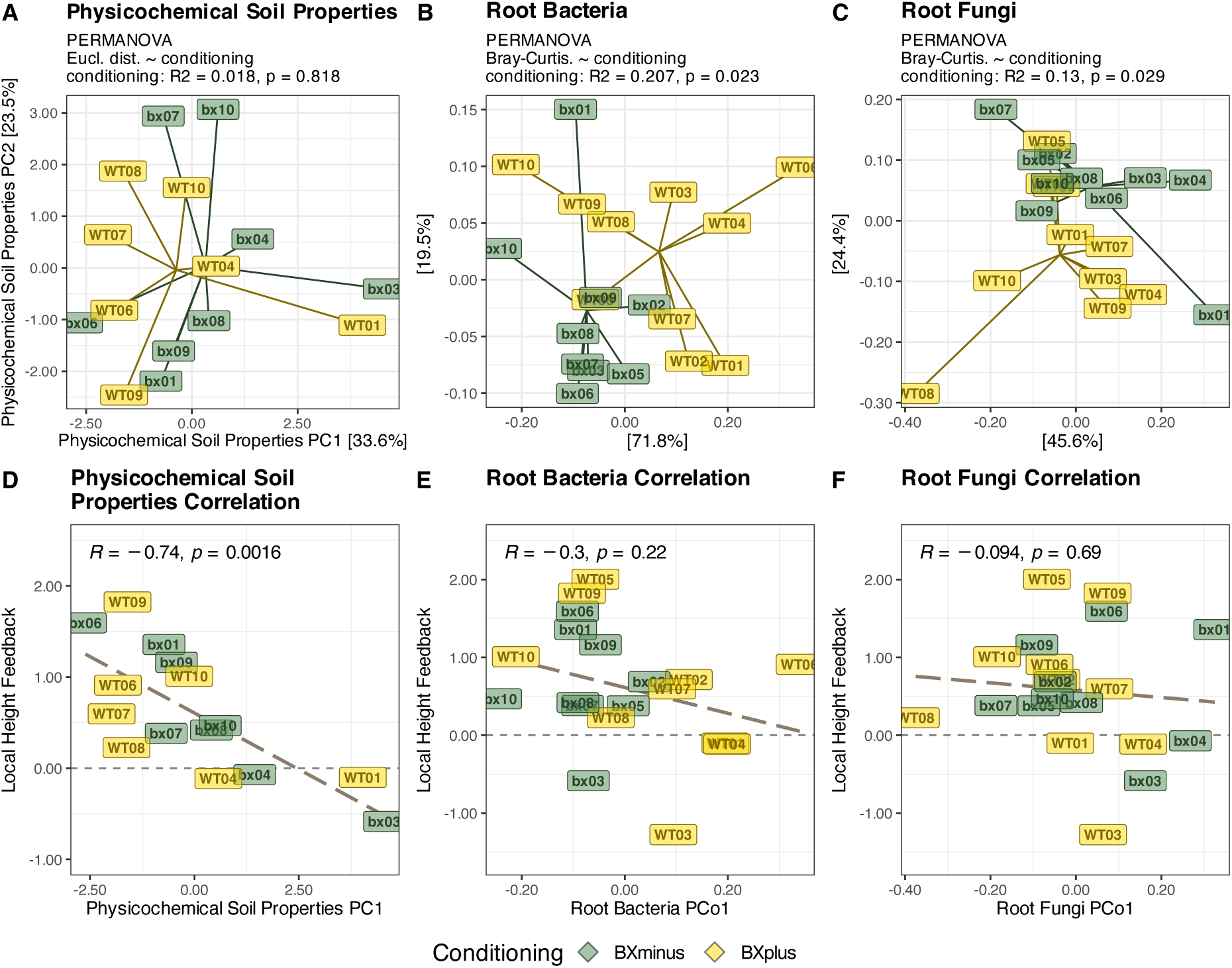
| Soil properties, maize microbial communities and their relationships to wheat’s local height feedbacks in Reckenholz. **(A)** Principal Component (PC) Analysis visualizes the variation in physicochemical soil properties (Euclidean distance, PC axes 1 and 2 with % of variation explained are shown) while Principal Coordinates (PCo) Analysis depicts variation among **(B)** bacterial and **(C)** fungal community profiles of maize roots at the end of the conditioning phase (Bray-Curtis distance, PCo axes 1 and 2 with % of variation explained are shown). The effects of conditioning were tested for soil physicochemical parameters and bacterial and fungal ASVs using permutational analysis of variance (PERMANOVA), with the model and corresponding R^2^– and p-values displayed above the plots. Next, we tested the interdependence of wheat feedbacks with the soil chemical or microbial properties. The axes with highest correlation coefficient, explaining at least 10% of variation, were chosen for display (see **Table S1**). The local height feedbacks correlated best with **(D)** the first PC axis of the physicochemical soil parameters, **(E)** the first PCo axis of bacterial communities, and **(F)** the first PCo of fungal communities. The corresponding Pearson’s correlation coefficients (R) and p-values are indicated at the top of each panel. Colors represent the conditioning treatments.

Next, we evaluated whether the wheat’s feedback response, i.e. the local height feedbacks, depends on the soil physicochemical or microbial parameters in each plot. For this, we conducted correlation analyses using the ordination axes of the physicochemical parameters and the microbial communities explaining at least 10% of variation (**Table S1**). Interestingly, the strength of wheat’s local height feedbacks correlated most strongly and significantly with the local soil parameters (physicochemical soil chemistry PC1; R = –0.74, *p* < 0.01, **Fig. 2D**) while this was not the case with the microbial communities (bacteria: PCo1, R = –0.30, *p* > 0.05, **Fig. 2E**; fungi: PCo1, R = –0.09, *p* > 0.05, **Fig. 2F**). Taken together, BX-producing and defective maize lines differentially condition the microbial communities but the local physicochemical soil parameters largely define the strength of the microbial feedbacks on wheat height.

### Dependence of wheat feedbacks on soil parameters is consistent across multiple fields

Next, we integrated the Reckenholz finding, that local soil heterogeneity links with varying feedback strength, to our earlier field experiments conducted at Posieux (Gfeller et al., 2023b) and Changins (Gfeller et al., 2023a; **Fig. 3A**). Analogous to above but for all three field experiments combined, we analysed the local wheat height feedbacks, physicochemical soil parameters, the microbial communities of maize plants and their interdependence. Wheat performance was significantly enhanced in BX_plus_ compared to BX_minus_ conditioned plots (*p* < 0.01, **Fig. 3B**). We noticed substantial variation in local height feedbacks within each field soil and examined also the soil parameters across the three fields.

**Figure 3.**
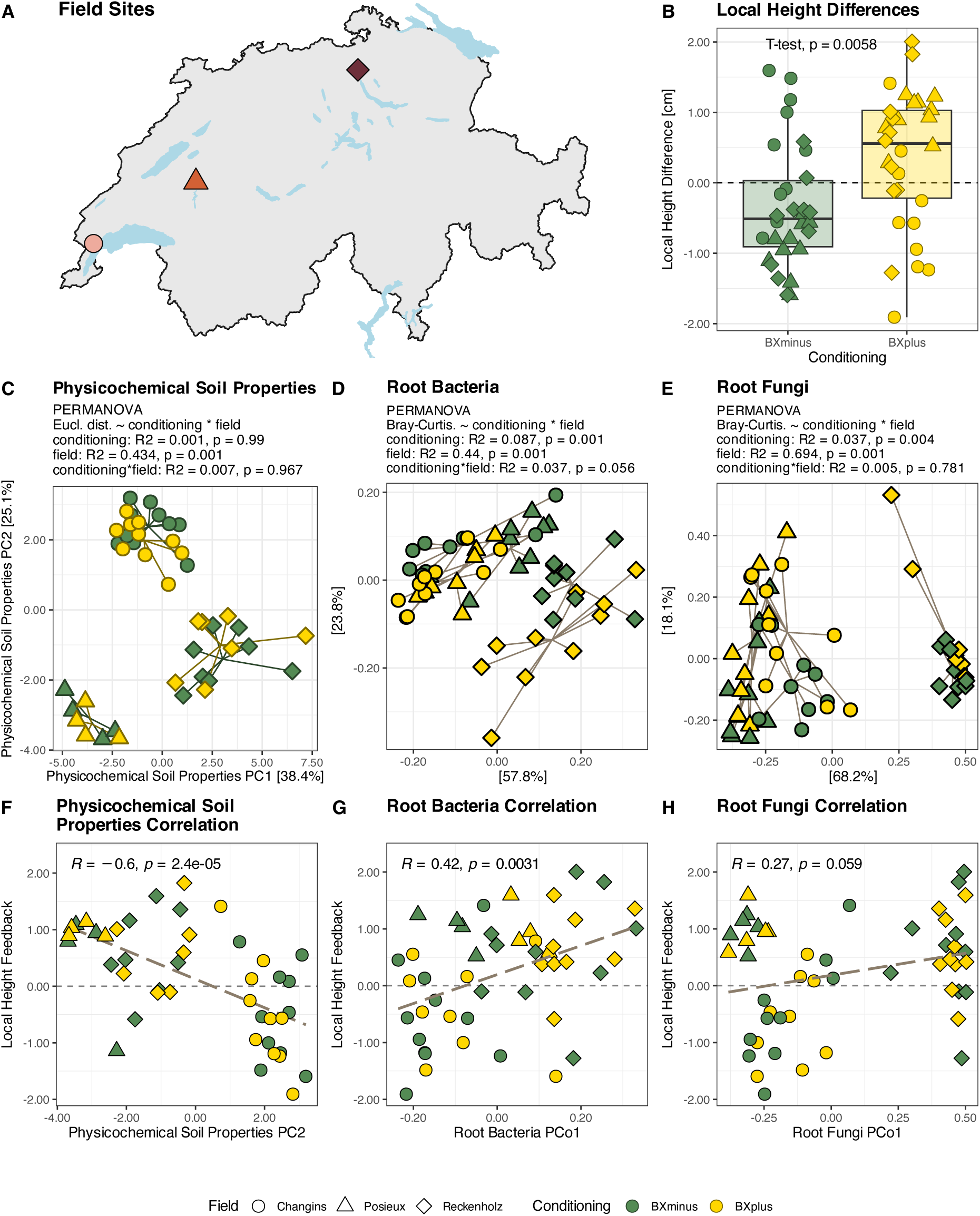
| Combinatory analysis of soil properties, maize microbial communities and their relationships to wheat’s local height feedbacks across three field experiments. **(A)** We have conduced plant-soil feedback experiments with wild-type and benzoxazinoid-deficient maize followed by wheat at the three field sites Changins, Posieux and Reckenholz in Switzerland. **(B)** We found no alteration in local height differences between wheat grown on soil conditioned with *bx1*– or WT-maize (t-test, p-value displayed in panel). **(C)** Principal Component (PC) Analysis was used to display the variation in physicochemical soil properties, as measured at the end of the conditioning phase (Euclidean distance, PC axes 1 and 2 with % of variation explained are shown). Principal Coordinates (PCo) analysis was used to visualize the variation among **(D)** bacterial and **(E)** fungal community profiles of maize roots at the end of the conditioning phase (Bray-Curtis distance, PCo axes 1 and 2 with % of variation explained are shown). The differences between sites, the effects of soil conditioning and their interaction were tested using permutational analysis of variance (PERMANOVA), with the model, R^2^– and p-values reported above the plots. Next, we tested the interdependence of wheat feedbacks with the soil chemical or microbial properties using correlation analysis. The axes with highest correlation coefficient, explaining at least 10% of variation, were chosen for display (see **Table S2**). The local height feedbacks correlated best with **(F)** the second PC axis of the physicochemical soil parameters and the first PCo axes of the **(G)** bacterial and **(H)** fungal communities. The corresponding Pearson’s correlation coefficients (R) and p-values are indicated at the top of each panel. Colors represent the conditioning treatments and shapes represent field site.

The physicochemical soil parameters differed significantly between the three field sites (PERMANOVA: R^2^ = 43.4% of the variance between sites, *p* < 0.01; **Fig. 3C**). This is not surprising with three different soil types: Reckenholz being silt-rich, Changins clayey, and Posieux sandy (**Fig. S3A**). It’s noteworthy that Changins soil contained more than twice as much plant-available (PA) iron compared to the other fields (**Fig. S3B**). Consistent with the findings above (**Fig. 2A**) and the earlier two field experiments (Gfeller et al., 2023a, 2023b), soil conditioning by BX exudation did not impact the physicochemical parameters (PERMANOVA: R^2^ = 0.001%, *p* > 0.05; **Fig. 3C**).

Next, we compared the microbial communities that maize plants leave as a legacy for wheat growth across the three field sites. Again, not surprising with three different soil types, the microbial communities significantly differed between the sites (PERMANOVA: bacteria: R^2^ = 44.0%, *p* < 0.01, **Fig. 3D**, fungi: R^2^ = 69.4%, *p* < 0.01; **Fig. 3E**). In contrast to the physicochemical soil parameters, the microbial communities were impacted by BX exudation: BXs caused an average shift of 8.7% in bacterial (*p* < 0.01) and 3.7% in fungal (*p* < 0.01) root communities. These findings are also consistent with the separated analyses of the field experiments (**Fig. 2BC**, and Gfeller et al., 2023a, 2023b).

Finally, we examined the interdependence of the varying strength of local feedbacks (**Fig. 3B**) with the variations in the soil properties across the three field sites. For this we correlated the local feedbacks with the ordination axes of the physicochemical or microbial communities (**Table S2**). The strength of local height feedbacks correlated most strongly with the physicochemical soil parameters (PC2, R = –0.6, *p* < 0.001; **Fig. 3F**) and to a lesser extent with the maize bacterial communities (PCo1, R = 0.42, *p* < 0.01, **Fig. 3G)** but not with fungal communities (PCo1, R = 0.27, *p* > 0.05, **Fig. 3H**). The combinatory analysis reinforced the conclusion of the Reckenholz site be valid across the three fields: BX-producing and defective maize lines differentially condition the microbial communities but the locally varying soil physicochemical parameters explain the variation in wheat’s feedback strength. This analysis further highlighted the additional involvement of local bacterial communities.

### Plant-available iron is the strongest predictor of local height feedbacks

To identify the explanatory parameter(s), we modelled the physicochemical soil measures and the abundant ASVs of the maize microbial communities relative to the local height feedbacks across the three field sites. We used repeated subsample-based LASSO regression with stability analysis (step 1) followed by stability-based filtering (step 2, see methods; **Fig. S4A**). A total of 63.6% of the variation in wheat height feedbacks could be explained by four parameters. The most reliable predictor was PA-iron, explaining 18.9% of variation and having a strong stability of 91%. Three weak-stable bacterial predictors explained together around 45% of variation and belong to the genera *Bacillus* (bASV17, explained variation: 14.1%, stability: 30%), *Pseudomonas* (bASV45, v: 13.6%, s: 33%) and *Mesorhizobium* (bASV697, v: 17.0%, s: 41%). No fungal predictor was found (**Fig. 4A**, **Fig. S4B**).

**Figure 4.**
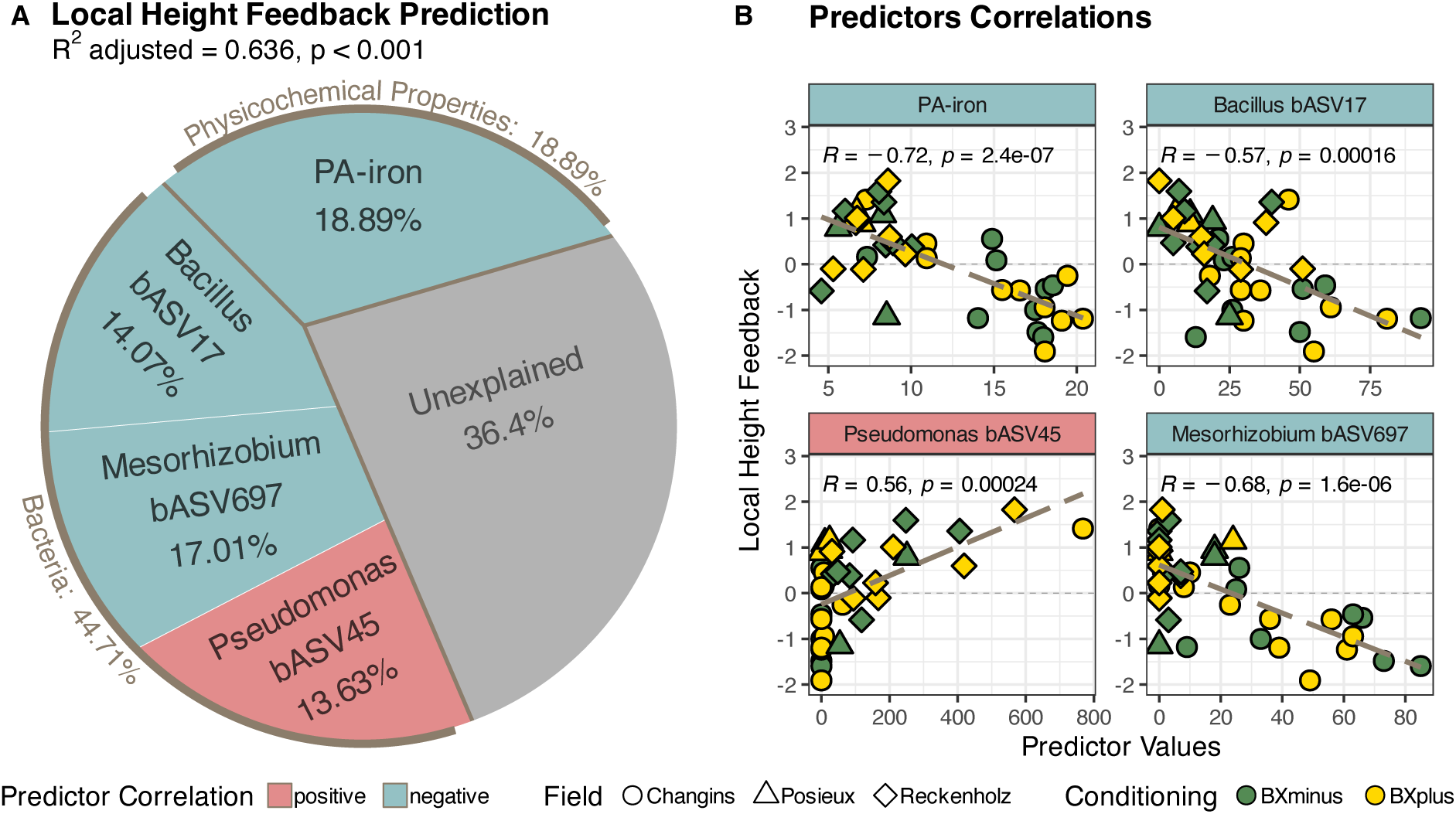
| Modelling local height feedback using physicochemical soil properties, bacterial ASVs and fungal ASVs. **(A)** Predictor selection for explaining local height feedback was done by a least absolute shrinkage and selection operator (LASSO) regression with a repeated subsample-based variable stability. The total model performance was adjusted by the number of selected predictors. Adjusted R^2^ and p-value is indicated above the plot. The Voronoi plot shows selected predictors. The proportion of explained variation based on adjusted R^2^ in local height feedback is labelled for each predictor and for predictor groups. **(B)** Local height feedback was correlated with each selected predictor. Pearson’s correlation coefficient (R) and p-value are labelled at the top of each panel. Colours represent predictor correlation directions and conditioning treatments. Shapes represent field sites.

PA-iron was negatively correlated with local height feedbacks (R = –0.72, *p* < 0.001; **Fig. 4B**) – i.e., the fewer iron there is available in soil, the more positive the local height feedback is on wheat. bASV17 and bASV697 were also negatively correlated with the local height feedback (bASV17: R = –0.57, *p* < 0.001; bASV697: R = –0.68, *p* < 0.001) while bASV45 showed a strong positive correlation (R = 0.56, *p* < 0.001). Interestingly, we also found that PA-iron concentrations, bASV17 and bASV697 abundances were strongly correlated with the soil BX concentrations in BX_plus_ conditioned soil samples, while this was not the case for bASV45 (**Fig. S5**; DIMBOA (PA-iron: R = 0.82, *p* < 0.001; bASV17: R = 0.73, *p* < 0.05, bASV697: R: 0.74, *p* > 0.001; bASV45: R = –0.36, *p* > 0.05) and DIMBOA-Glucose (PA-iron: R = 0.82, *p* < 0.001; bASV17: R = 0.73, *p* < 0.05, bASV697: R: 0.71, *p* > 0.05; bASV45: R = –0.34, *p* > 0.05)). Summarising, we identified one soil parameter, PA-iron, and three maize bacteria, of which PA-iron was the strongest and most reliable predictor to explain the variation in local height feedbacks of wheat.

### Low levels of PA-iron link with beneficial microbial feedbacks on plant growth

The finding that PA-iron in soil explained the strength of wheat feedbacks was reminiscent of the field experiment in Changins, where the chemical gradient in soil was particularly driven by high levels of PA-iron at one end of the field site (Gfeller et al., 2023a). From that field site in Changins, we regularly collected soil batches to study microbiome feedbacks on Arabidopsis (details in **Table S3**) and noticed that the strength of the feedbacks (measured as differential rosette area) also differed between soil batches (**Fig. S6**). Therefore, we reasoned that the levels of PA-iron might also explain the varying feedbacks on Arabidopsis growth. While the soil batches for Arabidopsis experiments were randomly collected without recording exact field coordinates, we had them analysed for their physicochemical soil parameters.

To probe this idea, we correlated the varying feedbacks on Arabidopsis growth with the different PA-iron concentrations in each soil batch. Consistent with the field work with wheat (R = –0.71, *p* < 0.001), we also found a significant negative correlation between the Arabidopsis growth feedbacks and PA-iron in soil (R = –0.5, *p* < 0.05; **Fig. 5A**). These correlations did not differ from each other (Fisher: z = 1.2; *p* = 0.231), suggesting that microbial feedbacks on wheat and Arabidopsis follow a similar relationship with soil PA-iron levels. Therefore, we saw support for the idea that soil PA-iron also explains the varying strength of Arabidopsis growth feedbacks in the different soil batches.

**Figure 5.**
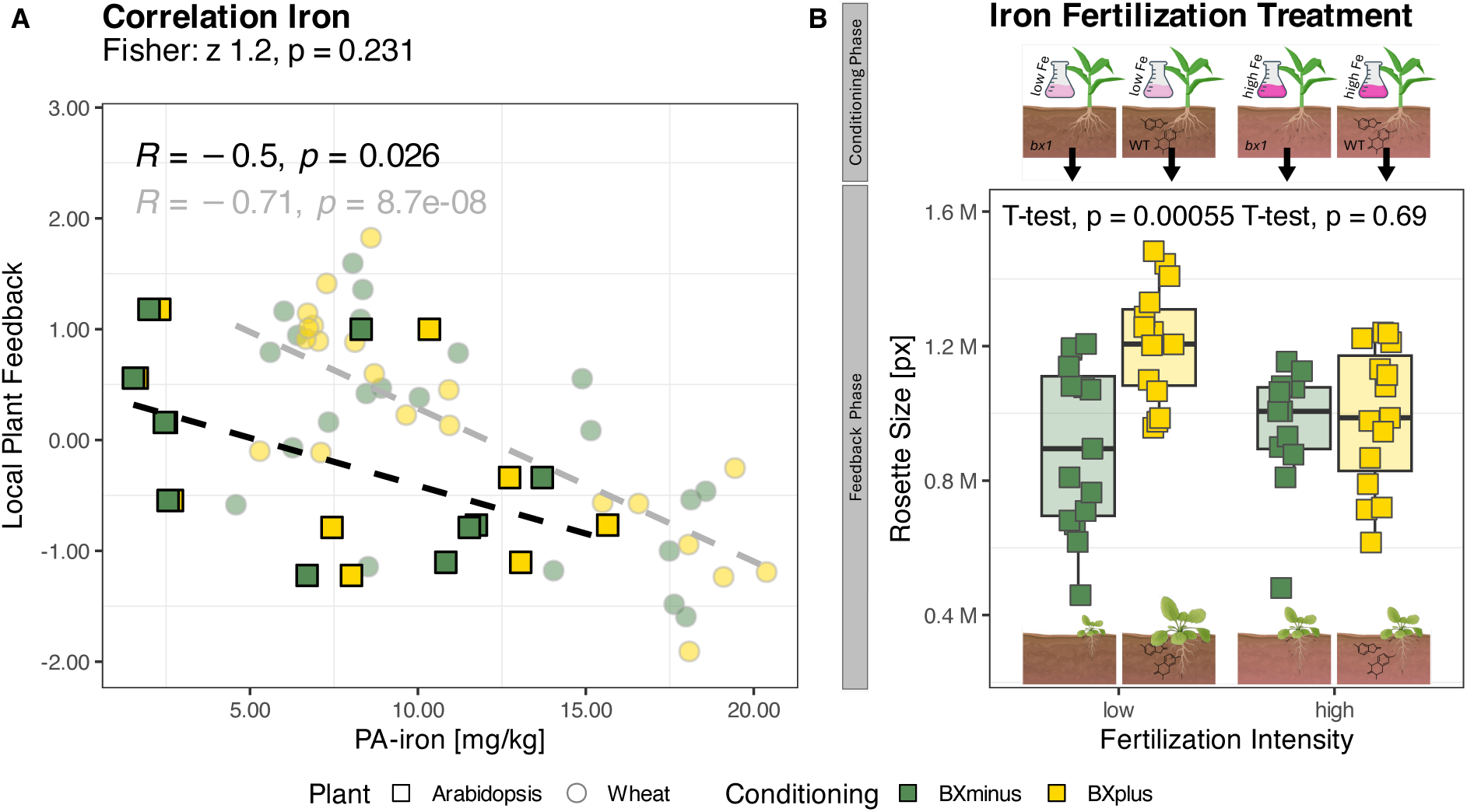
| Meta-Analysis of maize-Arabidopsis plant-soil feedback experiments. **(A)** Local rosette feedback of Arabidopsis, respectively local height feedback from wheat, was calculated and correlated with plant-available iron (PA-iron). Pearson’s correlation coefficient (R) and p-value are labelled at the top. With a Fisher test it was investigated for differences in the correlation coefficients between the two independent groups of Arabidopsis and wheat. Test statistic z and p-value are labelled above the plot. **(B)** At the end of the feedback phase, differences in rosette size of Arabidopsis grew on BX_plus_ and BX_minus_ conditioned soil were tested using a t-test. Colours represent data origins, local feedback directions and conditionings. Opacities and shapes refer to maize-wheat experiments and maize-Arabidopsis experiments.

Emerging from the wheat and Arabidopsis feedback findings, we hypothesised that increasing soil PA-iron levels would suppress the positive microbial feedbacks on plant growth. To test this, we conditioned Changins soil with wild-type and *bx1* maize plants at both high and low levels of iron fertilization and subsequently scored the microbial feedbacks on Arabidopsis growth (**Fig. 5B**, upper part). Consistent with this hypothesis, Arabidopsis grew smaller on BX_minus_ compared to BX_plus_ soil in low iron conditions (*p* < 0.001) while high iron supplementation during soil conditioning eliminated this positive feedback and Arabidopsis grew equally on BX_minus_ and BX_plus_ soils (*p* > 0.05; **Fig. 5B**). Hence, with this iron supplementation experiment, we could experimentally validate the correlative evidence for an inverse relationship between soil PA-iron levels and microbial feedbacks on plant growth (wheat, **Fig. 4B**; Arabidopsis, **Fig. 5A**).

## Discussion

It is well established that (i) plants change their surrounding soil microbiome by the release of root exudates, (ii) such conditioned microbiomes impact the performance of a new plant generation and (iii) that such ‘microbiome feedbacks’ depend much on the soil context. A key knowledge gap towards predictability is the identification of the underlying factors that explain the variation in strength and direction of soil microbiome feedbacks on plant growth (Cheng et al., 2024; van der Putten et al., 2016). Here, we reported a new field site where local soil heterogeneity explained the variation in microbiome feedbacks on wheat height. We used this data combined with data of previous field experiments to identify the factor(s) that explain the variability of microbial feedbacks using multivariate, correlation and modelling analyses. Across multiple field sites and extended to Arabidopsis, we found that soil levels of PA-iron largely explain the varying strength of the microbiome feedbacks. Below, we describe the emerging model and discuss its possible underlying mechanisms and the implications of soil heterogeneity.

### Model: positive feedbacks at low iron levels

Analysing the new field data from the Reckenholz site, alone as well as when combined with the Posieux and Changins sites, revealed consistency in two things: first, maize’s legacy microbiomes differed between wild-type and *bx1* plants (**Figs. 2B&C, 3D&E**) and second, the strength of their resulting feedbacks on wheat largely depended on the physicochemical soil parameters (**Figs. 2D, 3F**). PA-iron was the most important and most stable predictor to explain local height feedbacks of wheat (**Fig. 4**). We found this inverse relationship between soil iron and wheat feedbacks also to apply to Arabidopsis feedbacks where the strength varied with iron levels in the different soil batches collected from the Changins site (**Fig. 5A**). The following model emerges from taking all findings of this study together (**Fig. 6**): the microbial communities in BX_plus_ soils provoke positive feedbacks on wheat or Arabidopsis growth at low levels of PA-iron in soil while they cause no or even negative feedbacks at high iron levels. Testing this model with iron fertilisation revealed that the positive growth feedbacks of BX_plus_ microbiomes were lost at elevated soil iron levels (**Fig. 5B**). In conclusion, low levels of PA-iron link with beneficial feedbacks of the BX_plus_ microbiome on plant growth. Future work is needed to identify which microbes and how they promote plant growth under low iron conditions.

**Figure 6.**
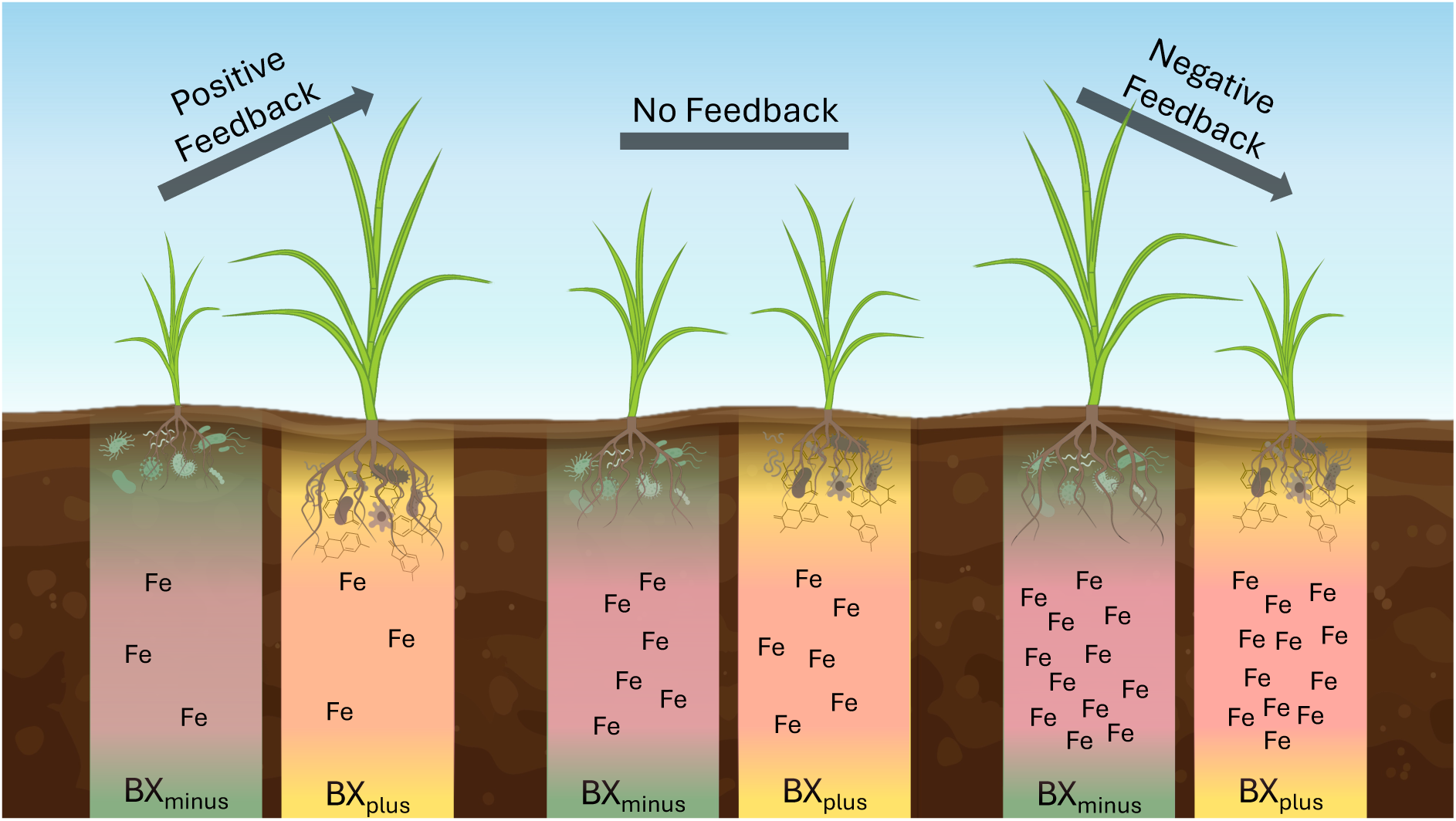
| Dependency of BX-mediated feedbacks on plant-available iron in wheat. Model illustrating how local plant-soil feedbacks in wheat, growing on BX_minus_ or BX_plus_ conditioned soil, are influenced by plant-available iron concentrations. At low plant-available iron levels, wheat performs better on BX_plus_ conditioned soil than on BX_minus_ conditioned soil. As plant-available iron increases, this performance difference diminishes. At high plant-available iron concentrations, the direction of the feedback reverses, and wheat grown on BX_minus_ conditioned soil outperforms wheat grown on BX_plus_ conditioned soil.

### Bacterial predictors

Next to PA-iron as highly stable predictor, three bacterial predictors have been detected (**Fig. S4B**). These were sequences assigned to a *Bacillus* (bASV17), a *Pseudomonas* (bASV45) and a *Mesorhizobium* bacterium (bASV697). *Bacillus* bASV17 and *Mesorhizobium* bASV697 were negatively correlated to the feedback on plant height while *Pseudomonas* showed bASV45 a positive correlation (**Figs. 4**). Previous studies support potential links between these genera, iron availability, and plant performance. For instance, *Bacillus altitudinis* WR10, an endophyte isolated from wheat, was shown to improve root and shoot growth of wheat under iron-deficient conditions by up-regulation of ferritin-related genes in roots (Sun et al., 2017). In field trials, the strain WR10 increased both grain iron content and agronomic traits such as kernel number per spike (Sun et al., 2021). Similarly, many Pseudomonas strains like GRP3A, PRS9 or RSP5 are well-known siderophore producers (Sharma and Johri, 2003). Inoculation with these strains under low iron conditions has shown increased Fe accumulation, higher seed germination rates and more biomass in cereals, such as maize. Additional these strains were also antagonists against phytopathogens such as *Colletotrichum dematium*, *Rhizoctonia solani* and *Sclerotium rolfsii* (Sah et al., 2017; Sharma and Johri, 2003). In contrast, *Mesorhizobium* species are primarily known for their nitrogen-fixing capabilities in legumes, and their role in iron mobilization in wheat remains less clear. A study by Verma et al. (2013) reported that a co-inoculation of a *Mesorhizobium sp.* BHURC03, a *Pseudomonas aeruginosa* strain BHUPSB02 and a *Bacillus megatorium* BHUPSB14 showed significant enhancement in growth and nutrient acquisition, such as phosphorus and iron, in chickpeas. Further investigation on these bacterial predictors is required and ideally bacterial strains matching the predictive *Bacillus* (bASV17), *Pseudomonas* (bASV45) and *Mesorhizobium* (bASV697) could be isolated for functional assays at varying levels of PA-iron in soil.

### BXs as iron chelators

The above-mentioned model is challenged by the known function of BXs to act as phytosiderophores chelating iron for plant uptake (Hu et al., 2018a). Therefore, it appears plausible that feedback plants may benefit from residual BX-iron complexes in soil. Interestingly, it has indeed been demonstrated that non-BXs-producing plants, such as oat and rice, can benefit from BX-iron chelates from the surrounding environment (Zhou et al., 2018). Naïvely, this would explain that Arabidopsis, also a non-producing species, might benefit from BX-iron complexes under low PA-iron conditions. However, there is a very clear finding that contradicts such a plausible explanation: Although not specifically quantifying BX-Fe complexes, we found very strong positive correlations between soil levels of PA-iron and both DIMBOA-Glu and DIMBOA levels in soil (**Fig. S5**). These two compounds are the most abundantly exuded BXs in maize (Hu et al., 2018b) and are the BX species that actually chelate iron (Hu et al. 2018, Science). The naïve explanation cannot work because the growth benefit is seen at low iron conditions, where the measured levels of soil DIMBOA-Glu and DIMBOA were also low. Taken together, it is unlikely that the phytosiderophore function of plant BXs would explain the growth benefit seen at low iron conditions. Instead, BXs might function to recruit a beneficial microbiome under low-iron conditions where possibly the plants benefit from either more growth-promoting microbes and/or microbes with dedicated siderophores. These latter possibilities require experimental testing e.g. by comparing bacterial isolates from BX_minus_ and BX_plus_ soils for their capacity to promote growth and to mobilise or solubilise PA-iron.

### Local soil conditions – essential co-variable in field studies

The dependency of microbial feedbacks on their soil environmental context is well recognised (Cheng et al., 2024; van der Putten et al., 2016). While varying feedback responses are commonly found comparing different field soils, they are less obvious and require higher numbers of replicate plots to detect them in a fine-grained manner within fields. Each of our field experiment was conducted with 10 replicated strips (Posieux) or plots (Changins, Reckenholz) per soil conditioning. At Changins, where the field has a characteristic physicochemical gradient in soil, the BX-induced microbial feedbacks significantly changed along the gradient, while the high variation did not allow to detect differences at field-scale (Gfeller et al., 2023a). Analogous at Reckenholz: wheat performance was unaffected (in statistical terms) when analysed at field-scale but significant differences were found between BX_minus_ and BX_plus_ soils when considering the *local* feedback (height, **Fig. 1C**; other phenotypes, **Fig. S2**). These examples highlight the importance of considering local soil heterogeneity for analysis and interpretation of field studies. In contrast to pot experiments, where field soil can be homogenised before filling, field studies require fine-grained analyses of the local soil parameters to account for spatial heterogeneity. We argue that the local soil physicochemical parameters present an essential co-variable for field studies to successfully disentangle the complex relationships between plant performance in response to the microbes in a given soil environment.

### Soil heterogeneity – essential for microbiome feedback experiments

Ironically, we had ignored the above conclusion for our experiments with Arabidopsis where we neglected local soil heterogeneity when sampling the different field soil batches. In the past, we did not pay attention *where* in the field we sampled soil for our pot experiments. Arabidopsis feedbacks were consistent within but inconsistent between different soil batches (**Fig. S6**, **Table S3**). Positive feedbacks were recorded for soil batches BS04, BS07 or BS09 while negative feedback was seen with the others. The strong relationship between soil PA-iron and feedback strength in wheat (**Fig. 4** and Gfeller et al., 2023a) prompted us – fortunately we had all batches analysed for their physicochemical soil parameters – to examine PA-iron as possible driver of the varying feedbacks on Arabidopsis growth. Importantly, the significant negative relationship between Arabidopsis feedbacks and PA-iron (**Fig. 5A**) was consistent with wheat and could be validated with a fertilisation experiment. Using a soil batch with low iron levels, the positive growth feedback was lost upon enhancing soil iron levels (**Fig. 5B**). In retrospect, we learned the important lesson that local soil heterogeneity, not only as co-variable for field studies, is also essential for microbiome feedback experiments with natural soil under controlled conditions.

### Conclusions

The initial observation that microbiome feedbacks are highly dependent on the soil physicochemical parameters could be narrowed down to PA-iron as the main factor. The key finding of this study is that the levels of PA-iron in soil largely define the strength of microbial feedbacks both on wheat and Arabidopsis. Low levels of PA-iron were needed for the beneficial soil microbiota (in this study, conditioned by BX-producing plants) to provoke positive growth feedbacks. In contrast, higher levels of PA-iron resulted in no– or even negative microbial feedbacks. An indirect conclusion of this finding is that fertilisation, i.e. enhancing PA-iron levels, is derogatory to beneficial microbial feedbacks. In simplified terms, when plants are nutritionally supported by fertilisers, they do not need or benefit from beneficial soil microbiomes. Thinking practically – e.g., about crop rotation at reduced fertiliser inputs – it will be important i) to understand the environmental factors (e.g., low iron conditions) and ii) to identify beneficial plant microbiomes (e.g., of BX producing maize plants) that then positively feedback on the next crop generation. It comes down to solving the context-dependency of microbial feedbacks, allowing then to take informed decisions for microbiome-assisted cropping practices that eventually enhance agricultural sustainability.

## Author Contributions

**Jan Waelchli:** Conceptualization; investigation; methodology; field experiment Reckenholz; data curation; statistics; visualization; writing. **Henry Janse van Rensburg**: Investigation, Arabidopsis experiments. **Katja Stengele**: Arabidopsis experiments. **Viola D’Adda**: Arabidopsis experiments. **Selma Cadot:** field experiment Reckenholz. **Vero Caggìa**: field experiment Reckenholz. **Valentin Gfeller**: new data from Changins and Posieux and integration. **Klaus Schlaeppi**: supervision; conceptualization; investigation; methodology; writing.

## Acknowledgements

A special thanks to Friedrich Käser, Andreas Kägi, Daniel Fuchs and Beatrix Lanzini from Agroscope Reckenholz for access to the field and their support to make this 2-year field experiment possible. Further, we thank Profs. Matthias Erb and Christelle Robert for providing wild-type W22 and *bx1*(W22) maize seeds, Markus Funk and Dr. Lea Stauber for field assistance, Ellen Radon for the microbiome library preparation in the laboratory and Dr. Pamela Nicholson and her team at the Next Generation Sequencing Platform at the University of Bern (ngs.unibe.ch) for excellent help with sequencing. This work was mainly supported by fundings of the University of Basel and partly the University of Bern (Interfaculty Research Collaboration “One Health” to K.S.) and the Swiss National Science Foundation (No. 189249 to K.S.).

## Ethic statement

This research did not include human participants, nor animal experimentation or handling of biological material under protection by the Nagoya protocol. The authors confirm that the work is original, the data accurate and not manipulated and that all sources have been cited.

