## Supplementary File 1 for "Soil iron drives beneficial maize microbiome feedbacks in rotations with wheat"

Index

- Supplementary figures
- Supplementary tables

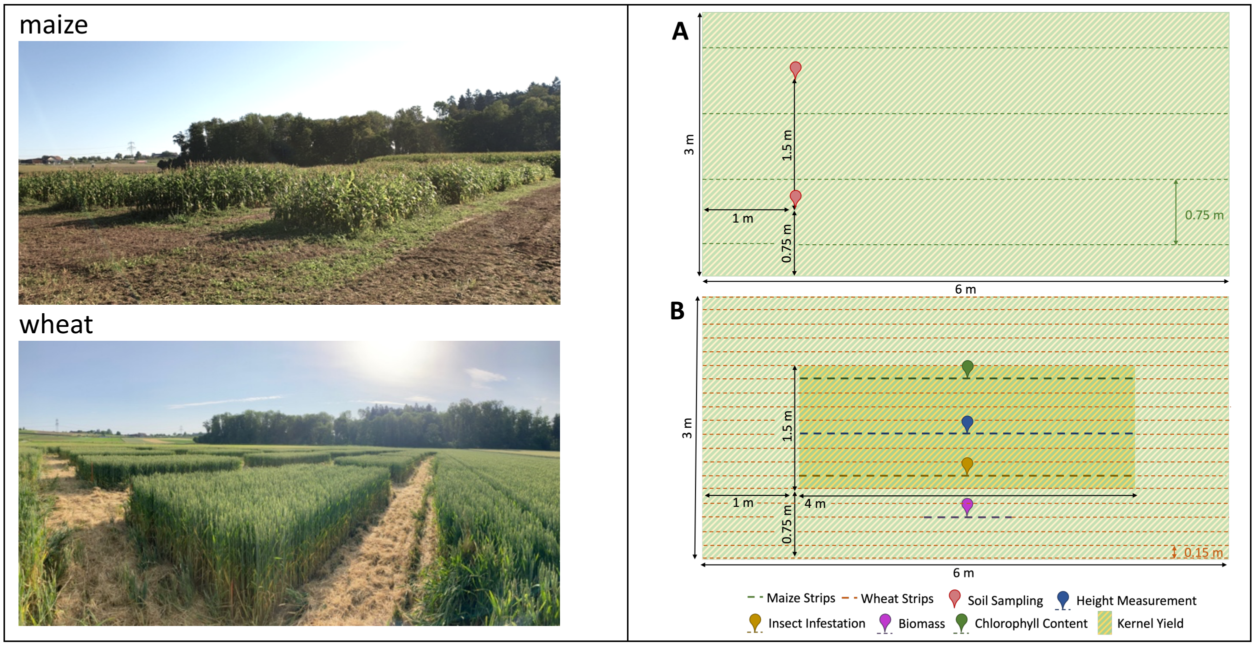


**Figure S1 | Experimental setup per field plot. (A)** For conditioning, maize was grown in strips with 75 cm distance. At the end of the conditioning phase, two soil samples per plot were taken marked by red pins. **(B)** During feedback phase wheat was grown in strips with 15 cm between them. Wheat height was measured from 10 randomly selected plants on the blue labelled line, *Oulema melanopus* larvae were counted from 10 randomly selected plants on the yellow line and the chlorophyll content were measured from 10 randomly selected plants on the green line. To measure the biomass, 1 meter of wheat was harvested, dried and weighted. After wheat has produced kernels, the core of each plot was harvested and the kernel weight was determined.

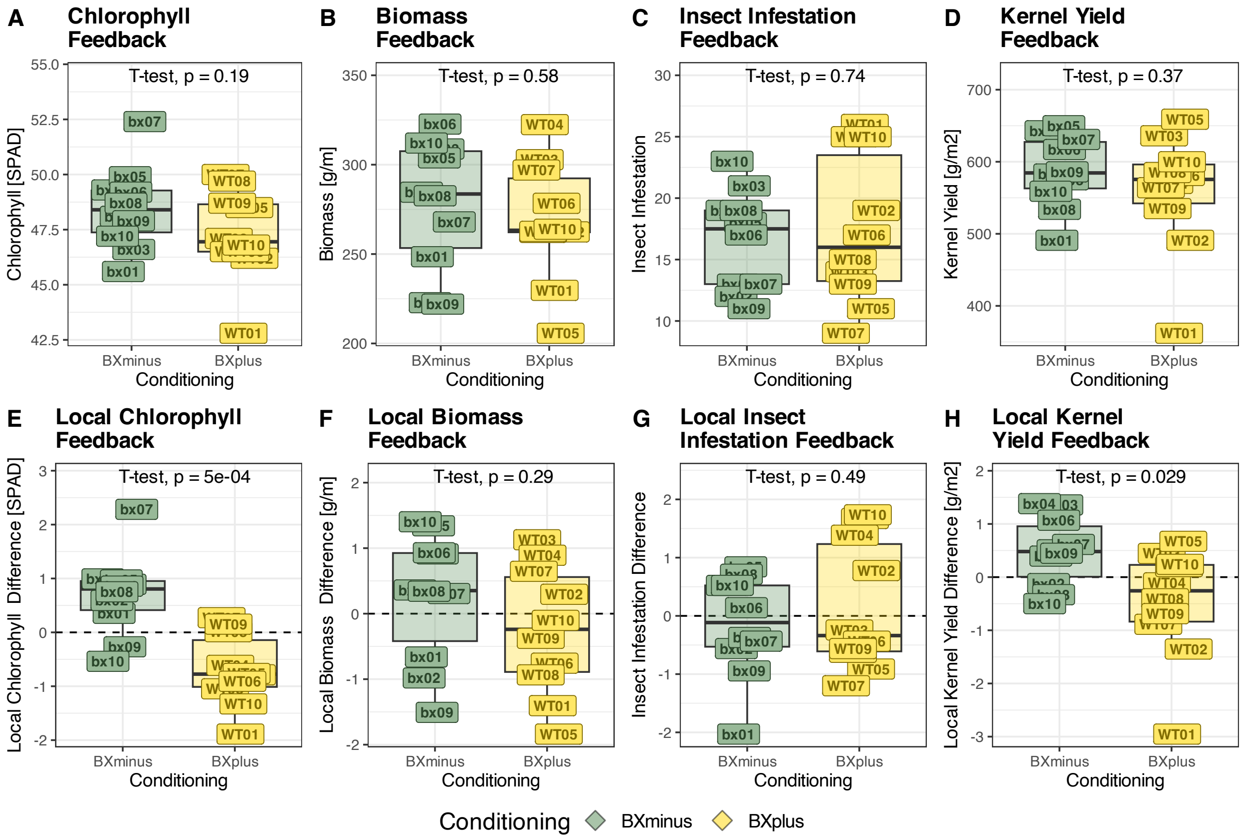


**Figure S2 | Wheat phenotyping.** Wheat phenotypes were measured towards the end of the feedback phase. The plant height is shown in **Fig. 1C** while here we show the measured chlorophyll content **(A)**, the dried plant biomass **(B)**, the number of *Oulema melanopus* larvae on the two newest tillers **(C)** and the kernel yield **(D)**. For none of them we find a difference between wheat growing on $BX_{plus}$ or $BX_{minus}$ conditioned soil. Reducing the spatial variation by calculation local feedbacks (see methods) shows differences for the local chlorophyll content **(E)** and the local kernel yield **(H)** but not for the local biomass **(F)** and the local insect infestation **(G)** (t-test, p-value displayed in panel).

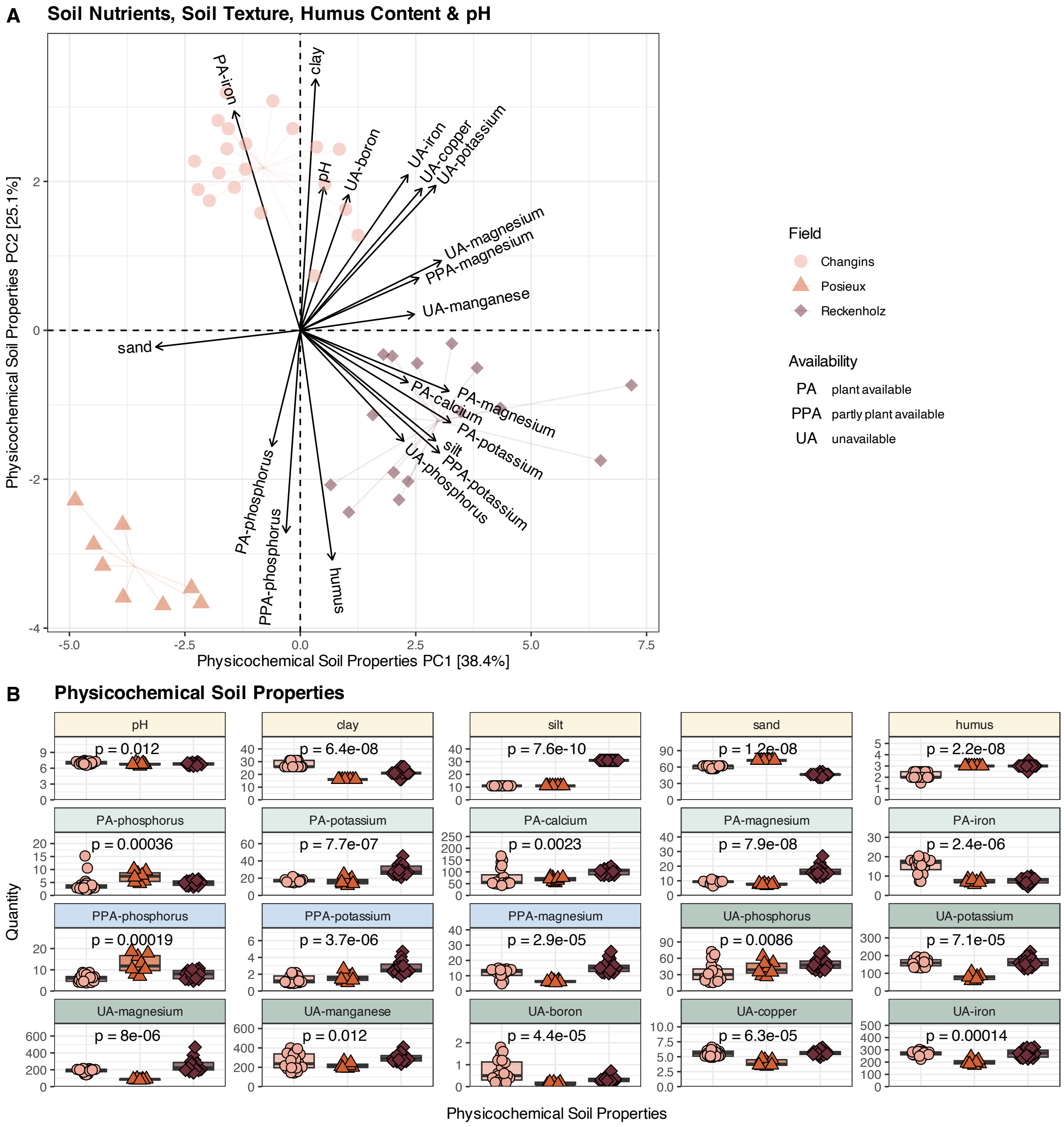


**Figure S3 | Physicochemical soil properties from all maize-wheat plant-soil feedback experiments.** Maize-wheat plant soil-feedback experiments were performed in Changins, Posieux and Reckenholz. **(A)** PCA show Euclidean distances between measured physicochemical soil parameters at the end of the conditioning phase with the corresponding loadings indicating the contribution of physicochemical soil properties. **(B)** The quantities of each soil property are tested for differences among the different field sites with an Analysis of Variance (ANOVA). P-values are corrected for multiple testing by the Bonferroni-Holm method. Soil properties are plant-available nutrients (PA), partly plant-available nutrients (PPA), unavailable nutrients (UA), soil texture, humus content and pH. Colours and shapes represent field sites.

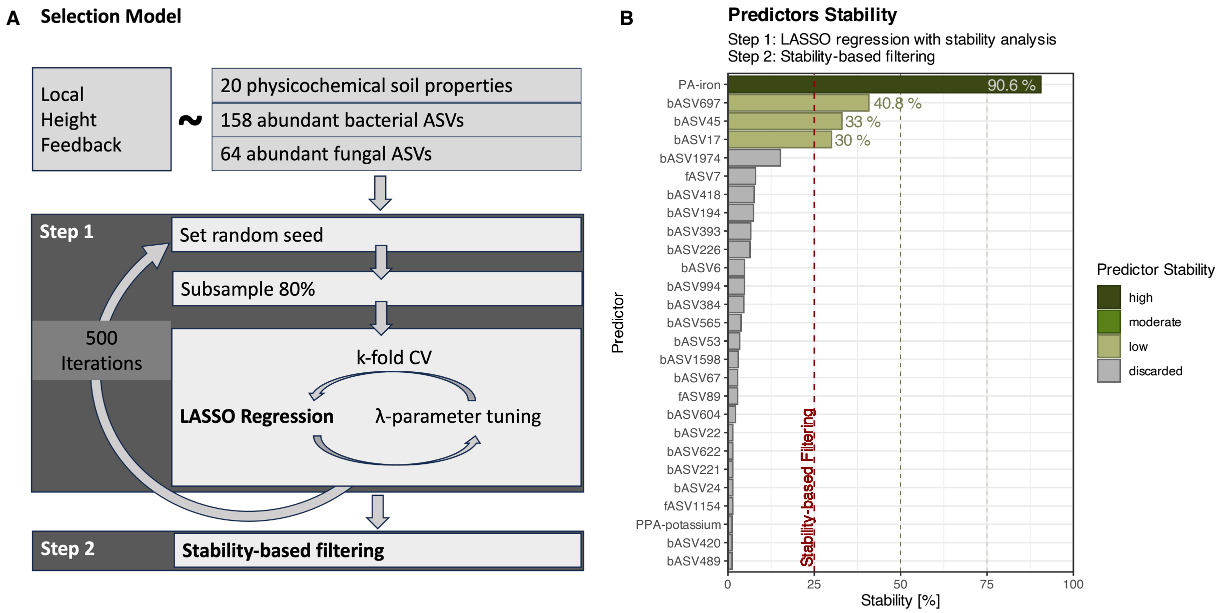


**Figure S4 | Three-Step Model Selection Approach. (A)** The local height feedback was modeled against 20 physicochemical soil properties and all bacterial and fungal ASVs with a minimum occurrence in at least 4 samples and a minimal total abundance of 5% (bacteria: 158 ASV, fungi: 64 ASVs). A least absolute shrinkage and selection operator (LASSO) regression approach was chosen with repeated subsample-based variable stability analysis. Step 1: For each of the 500 iterations a random seed was set, 80% of the samples were randomly selected as input and k-fold Cross-Validation was used to tune the λ-parameter. Step 2: Predictors with a lower selection frequency than 25% were considered as unstable and were discarded. **(B)** The plot shows the selection frequency of all predictors that have been identified by all LASSO regression with a frequency of at least 1%. Predictors were categorized by their selection stability: low (25–50%), moderate (50–75%) or high (75–100%).

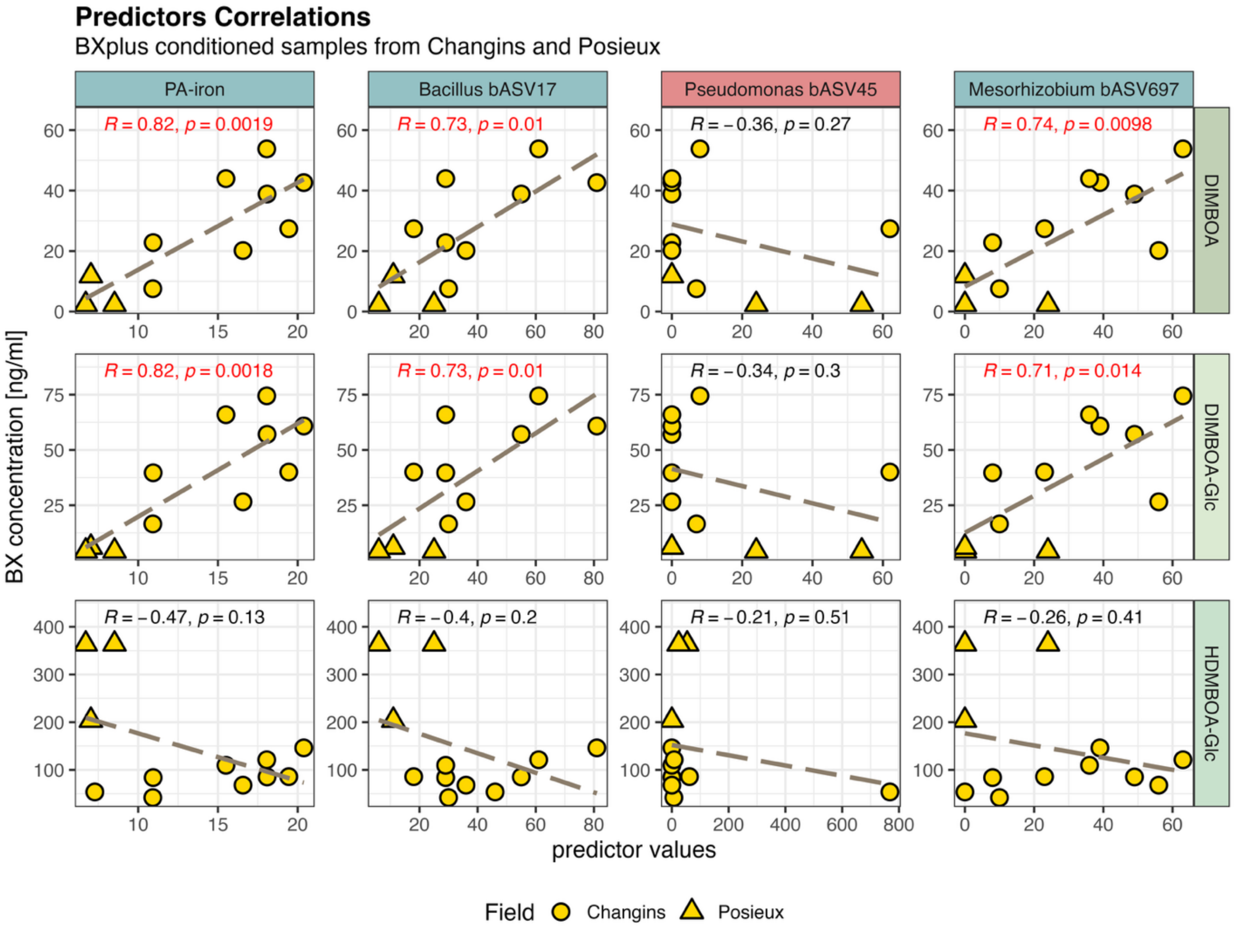


**Figure S5 | Predictors – Benzoxazinoids Correlation.** Benzoxazinoid-concentration has been measured on all $BX_{plus}$ conditioned plots in Changins and Posieux, but not in Reckenholz. Selected predictors were correlated with the 3 most exuded BX. Pearson's correlation coefficient (R) and Bonferroni-Holm adjusted p-value are labelled at the top. Significant p-values are marked in red. Shapes represent field sites. Colours of vertical facet titles represent the direction of correlation between predictor and local height feedback.

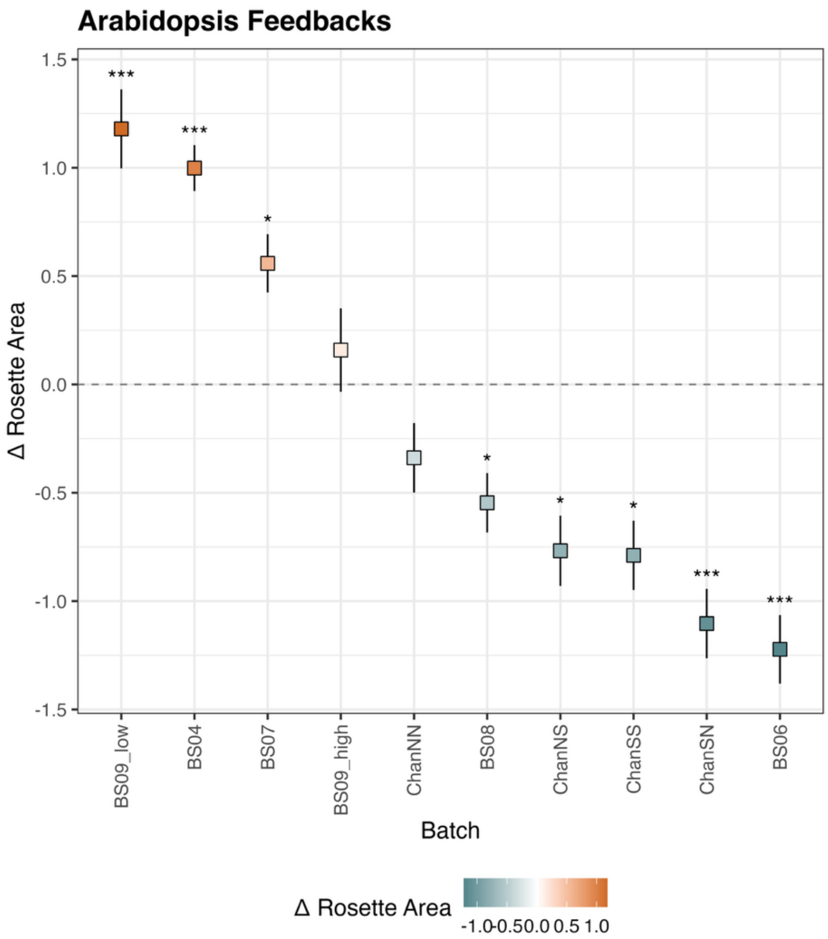


**Figure S6 | Arabidopsis: variation in feedback magnitude.** Rosette area feedbacks of Arabidopsis on $BX_{plus}$ or $BX_{minus}$ conditioned soils from different soil batches. Data were z-score normalised per experiment. Points represent mean differences +- standard errors. Positive values indicate Arabidopsis growing bigger on $BX_{plus}$ conditioning soil while Arabidopsis with negative values grow better on $BX_{minus}$ conditioned soil. For details on the calculation of local rosette area feedbacks, refer to the Materials and Methods section. A t-test per replicate was performed to determine differences between feedbacks on $BX_{plus}$ and $BX_{minus}$ conditioned soils. Asterisks mark significances (*<0.05, **<0.01, ***<0.001).

**Table S1: Correlation Reckenholz:** Correlation analysis of wheat's local height feedback to all principal component and coordinate axes of the soil physicochemical properties and microbial community data explaining at least 10% of variation. The $R^2$ values were derived from PCA and PCoA, respectively. The correlation coefficient rho (R-value) and p-value and were calculated with a Pearson correlation test. The axes with highest correlation coefficient (bold) were chosen for display in **Figure 2**.

| Reckenholz field site | | | |
|---|---|---|---|
| **Correlation Local Height Feedback to** | **$R^2$ value** | **R value** | **p-value** |
| **Physicochemical Soil Chemistry PC1** | **33.58%** | **-0.741** | **0.0016** |
| Physicochemical Soil Chemistry PC2 | 23.55% | -0.412 | 0.1272 |
| Physicochemical Soil Chemistry PC3 | 13.10% | -0.063 | 0.8238 |
| **Root Bacteria PCo1** | **71.82%** | **-0.297** | **0.2166** |
| Root Bacteria PCo2 | 19.46% | 0.102 | 0.6790 |
| **Root Fungi PCo1** | **45.65%** | **-0.094** | **0.6927** |
| Root Fungi PCo2 | 24.36% | -0.060 | 0.8010 |
| Root Fungi PCo3 | 14.53% | -0.069 | 0.7732 |

**Table S2: Correlation all field sites:** Correlation of local height feedbacks from Changins, Posieux and Reckenholz samples to all principal components of physicochemical properties or all principal coordinates of microbial communities explaining at least 10% of variation. The $R^2$ values were derived from PCA and PCoA, respectively. R-value and p-value were valuated with a person correlation test. Rows in bold label the strongest correlation axes used in figure 3.

| All field sites | | | |
|---|---|---|---|
| **Correlation Local Height Feedback to** | **$R^2$ value** | **R value** | **p value** |
| Physicochemical Soil Chemistry PC1 | 38.37% | 0.177 | 0.2559 |
| **Physicochemical Soil Chemistry PC2** | **25.06%** | **-0.597** | **2.40E-05** |
| Physicochemical Soil Chemistry PC3 | 16.56% | 0.317 | 0.0381 |
| **Root Bacteria PCo1** | **57.81%** | **0.423** | **0.0031** |
| Root Bacteria PCo2 | 23.76% | -0.118 | 0.4279 |
| **Root Fungi PCo1** | **68.24%** | **0.272** | **0.0589** |
| Root Fungi PCo2 | 18.08% | -0.186 | 0.2006 |

**Table S3: Collected soil batches to study microbiome feedbacks on Arabidopsis:** Multiple soil batches were collected on the field site in Changins. The soil was conditioned with WT and bx1 maize and the feedbacks on Arabidopsis were measured.

| Arabidopsis Datasets | | | | | | | | |
|---|---|---|---|---|---|---|---|---|
| | | | Conditioning Phase (Maize) | | | Feedback Phase (At) | | |
| **Batch** | **Nr of Exp** | **Availability** | **Growth (w)** | **Type** | **Fertilization** | **Fertilization (d)** | **Growth (d)** | **Type** |
| BS04 | 2 | (Janse van Rensburg et al., 2025) | 12 | pot | weekly (4w low[1]/8w high[2]) | 14[3], 21[3] | 35 | pot |
| BS04 | 1 | this study | 12 | pot | weekly (4w low[1]/8w high[2]) | 23[4] | 40 | pot |
| BS06 | 1 | this study | 12 | pot | weekly (4w low[1]/8w high[2]) | 15[4], 22[4] | 49 | pot |
| BS07 | 1 | (Janse van Rensburg et al., 2025) | 12 | pot | weekly (4w low[1]/8w high[2]) | 14[3], 21[3] | 35 | pot |
| BS07 | 1 | (Stengele et al., 2024) | 12 | pot | weekly (4w low[1]/8w high[2]) | - | 42 | pot |
| BS08 | 1 | this study | 12 | pot | weekly (4w low[1]/8w high[2]) | 14[3], 21[3] | 35 | pot |
| BS08 | 1 | this study | 12 | pot | weekly (4w low[1]/8w high[2]) | 23[4], 37[4] | 42 | pot |
| BS09_high | 1 | (Stengele et al., 2024) | 12 | pot | weekly (4w low[1]/8w high[2]) | 14[3], 21[3] | 35 | pot |
| BS09_low | 1 | (Stengele et al., 2024) | 12 | pot | weekly (12w low[1]) | 14[3], 21[3] | 35 | pot |
| ChanNN | 1 | this study | 12 | field | conventional farming practice | 15[4], 22[4] | AUC[5] | pot |
| ChanNS | 1 | this study | 12 | field | conventional farming practice | 15[4], 22[4] | AUC[5] | pot |
| ChanSN | 1 | this study | 12 | field | conventional farming practice | 15[4], 22[4] | AUC[5] | pot |
| ChanSS | 1 | this study | 12 | field | conventional farming practice | 15[4], 22[4] | AUC[5] | pot |

[1] Low iron: 100 mL of 0.2% Plantaaktiv Typ K, 0.001% Sequestrene Rapid

[2] High iron: 200 mL of 0.2% Plantaaktiv Typ K, 0.02% Sequestrene Rapid. All was applied once a week.

[3] Fertilization by watering with 10mL of 1/3 half strength Hoagland solution diluted in tap water

[4] Fertilization by watering with 5mL undiluted half strength Hoagland solution

[5]AUC stands for the area under the curve of weekly rosette area measurements from 3 to 8 weeks of growth
